# FMRP association with and regulation of Fragile X granules exhibit circuit-dependent requirements for the KH2 RNA binding domain

**DOI:** 10.1101/462168

**Authors:** Ellen C. Gingrich, Katherine A. Shepard, Molly E. Mitchell, Kirsty Sawicka, Jennifer C. Darnell, Michael R. Akins

## Abstract

The localization and translation of mRNAs is controlled by a diverse array of ribonucleoprotein particles (RNPs), multimolecular complexes containing mRNAs and RNA binding proteins. Fragile X granules (FXGs) are a family of RNPs that exemplify the diversity of RNA granules in the mammalian nervous system. FXGs are found in a conserved subset of neurons, where they localize exclusively to the axonal compartment. Notably, the specific RNA binding proteins and mRNAs found in FXGs depend on brain circuit and neuron type, with all forebrain FXGs containing Fragile X mental retardation protein (FMRP), the protein mutated in the human autism-related disorder Fragile X syndrome. FMRP negatively regulates FXG abundance but is not required for their association with ribosomes or mRNA. To better understand the circuit-dependent mechanisms whereby FMRP associates with and regulates FXGs, we asked how a disease-causing point mutation, I304N, in the KH2 RNA binding domain of FMRP affects these granules in two brain regions – cortex and hippocampus. We found that FMRP^I304N^ had a reduced association with FXGs, as it was absent from approximately half of FXGs in cortex and nearly all FXGs in hippocampus. FXG abundance correlated with the number of FMRP-containing FXGs, suggesting that FMRP regulates FXG abundance by KH2-independent mechanisms that occur locally within the granules. Together, these findings illustrate that cell type-dependent mechanisms guide the assembly of similar RNA granules. Further, point mutations in RNA granule components may lead to cell type-dependent phenotypes that produce atypical forms of disorders that normally arise from more severe mutations.

## Introduction

Fragile X syndrome (FXS), the most common inherited form of autism and intellectual disability, is caused by mutations in the *FMR1* gene encoding FMRP (Fragile X mental retardation protein) (Penagarikano et al., 2007). FMRP is a multifunctional protein that binds RNA in large multi-protein complexes to regulate protein synthesis and neuronal excitability in a wide range of developmental and cellular contexts (Bassell and Warren, 2008; Darnell and Klann, 2013). FMRP comprises multiple domains that mediate interactions leading to the assembly of a variety of FMRP-containing ribonucleoprotein particles (RNPs). These RNPs can differ in their abundance, RNA binding protein composition, and RNA association depending upon developmental stage, neuronal cell type, and subcellular compartment, consistent with the broad array of functions that FMRP carries out across the nervous system. This variety suggests that different mechanisms guide the formation of FMRP-containing RNPs in different parts of the nervous system.

Fragile X granules (FXGs) are a class of RNPs that exemplify how cellular context influences the formation and composition of FMRP-containing complexes (Akins et al., 2017; Christie et al., 2009; Chyung et al., 2018). FXGs are only found in axons and are not seen in somata or dendrites, despite the presence of the constituent proteins and mRNAs in these other cellular compartments (Akins et al., 2012, 2017; Christie et al., 2009). Moreover, FXGs are only found in a specific subset of neuronal populations that is stereotyped across individuals and conserved across mammalian species (Akins et al., 2012, 2017; Christie et al., 2009). Notably, the mechanisms governing FXG expression are cell type-dependent as FXGs are homogeneous within each neuronal population but vary across populations in their developmental window of expression, their protein composition, and the mRNAs with which they associate (Akins et al., 2017; Christie et al., 2009; Chyung et al., 2018). All forebrain FXGs contain FMRP along with the Fragile X related (FXR) protein FXR2P while circuit-selective subsets contain the FXR protein FXR1P. The remarkable spatiotemporal specificity of FXG expression and composition likely reflect a complex web of mechanisms that control their formation.

In past studies, we have used knockout mice to ask how the different FXR proteins (FMRP, FXR1P and FXR2P) regulate FXG expression. FXGs require FXR2P as brains of *Fxr2* null mice lack these granules (Christie et al., 2009). In contrast, neither FMRP nor FXR1P is required as even in wild type brains there are FXG populations that lack one or both of these proteins (Chyung et al., 2018). Instead, an increase in FXG abundance in *Fmr1* null mice reveals that FMRP regulates FXG number and the duration of their expression but does not regulate the circuits in which they are found or their structure (Akins et al., 2012, 2017; Christie et al., 2009; Chyung et al., 2018). Wild type FXGs have a characteristic substructure with the protein and RNA components arranged in overlapping but distinct subdomains. Loss of FMRP in *Fmr1* null brains does not impact the subgranular arrangement of the other FXG components, suggesting that these other components form the FXG core and FMRP affiliates with the granules by binding to one or more of them. These knockout studies could not address the mechanisms by which FMRP associates with FXGs nor could they identify how FMRP regulates FXG abundance.

Addressing these questions requires FMRP mutants in which select functions are perturbed while sparing others. A single amino acid substitution (I304N) of isoleucine to asparagine at FMRP residue 304 in the second hnRNP K homology (KH2) RNA binding domain was initially identified as a de novo mutation in a patient with particularly severe FXS (De Boulle et al., 1993). The I304N mutation greatly reduces the affinity of FMRP for kissing complex RNA and ribosomes and abrogates the ability of this protein to serve as a negative translational regulator (Feng et al., 1997; Laggerbauer et al., 2001; Zang et al., 2009). The mutant protein also does not bind to itself or to some other proteins including the Tudor domain containing protein Tdrd3 and the p21-activated kinase PAK1 (Hayashi et al., 2007; Laggerbauer et al., 2001; Linder et al., 2008; Say et al., 2010; Zang et al., 2009). However, FMRP^I304N^ retains the ability to assemble into large multimolecular complexes as well as to bind to G-quartet RNAs along with many proteins including FXR1P and FXR2P (Feng et al., 1997; Laggerbauer et al., 2001; Zang et al., 2009). FMRP^I304N^ may thus reveal whether KH2-mediated interactions are critical for FMRP association with FXGs.

We addressed this question in brains from *Fmr1*^I304N^ mice. Since FXGs differ in their composition across brain regions, mutations that impact FMRP interactions may differentially impact FXGs depending on cellular and developmental context and thereby reveal mechanisms that drive FXG formation in these various contexts. To address this possibility, we asked whether there are mechanistic differences in how FMRP associates with FXGs in two forebrain regions, cortex and hippocampus, that differ in FXG protein and mRNA composition. Indeed, we found that the I304N substitution attenuates FMRP association with FXGs in a circuit-dependent manner with FMRP present in most cortical FXGs but absent from nearly all hippocampal FXGs. Interestingly, FMRP regulation of FXG abundance depended on the presence of FMRP in the granules but not on whether FMRP could bind to RNA. FMRP thus limits FXG abundance by interactions that occur within FXGs but do not require FMRP regulation of protein synthesis. These findings suggest that FMRP associates with FXGs via KH2-dependent and KH2-independent interactions but that the relative contributions of these mechanisms depend on the cellular context in which they occur.

## Materials and Methods

### Animals

All work with animals was performed in accordance with protocols approved by the Institutional Animal Care and Use Committees of Drexel and Rockefeller Universities. Since the *Fmr1* allele is on the X chromosome, we used two parallel breeding schemes: one with dams heterozygous for the *Fmr1* null allele, which produced wild type and *Fmr1* null male littermates. In the second, the breeding mice were homozygous for the *Fmr1*^I304N^ allele; males were used from the resulting litters to minimize potential confounds of sex on FXGs, although our past studies have revealed no effect of litter or sex on FXG abundance or composition. Mice were deeply anesthetized before intracardiac perfusion with room temperature PBS (10 mM phosphate, pH7.4, 138 mM NaCl, 2.7 mM KCl) containing 1 U/mL heparin followed by perfusion with ice-cold PBS containing 4% paraformaldehyde. After perfusion, animals were decapitated, and intact brains removed, postfixed overnight in the perfusate and washed in PBS. Brains were then transferred to PBS with 30% sucrose until they sank and then embedded in OCT medium with rapid freezing at -80°C. A Leica CM3050 S cryostat was used to section 40 μm free-floating sagittal sections. Sections were stored in PBS and 0.02% NaN_3_ at 4°C.

### Immunofluorescence

Tissue was washed in PBS (10 mM phosphate, pH 7.4; 150 mM NaCl). To improve antibody access to epitopes, tissue sections were heated in 0.01M sodium citrate (pH 6.0) for 30 min at 75°C. Tissue was then treated with blocking solution [PBST (10 mM phosphate buffer, pH 7.4, and 0.3% Triton X-100) and 1% blocking reagent (Roche)] for 30 minutes. Sections were then treated with blocking solution plus primary antibody overnight, washed for 5 min with PBST, and then incubated for one hour in blocking solution plus appropriate secondary antibodies (1:1000; Life Technologies) conjugated to Alexa fluors. Sections were washed in PBST and then slide-mounted in NPG mounting medium (4% n-propylgallate, 85% glycerol, 10 mM phosphate buffer, pH 7.4). Monoclonal primary antibodies were generated in house from hybridoma supernatants and included 1G2 anti-FXR2P (1:300)(Zang et al., 2009), A42 anti-FXR2P (1:500)(Zhang et al., 1995), Y10b anti-rRNA (1:1000)(Garden et al., 1994; Lerner et al., 1981), and 2F5-1 anti-FMRP (1:500)(Christie et al., 2009; Gabel et al., 2004).

### In situ hybridization

This protocol was performed as previously described (Akins et al., 2017). In brief, fresh frozen brains were sectioned at 20 μm, mounted on slides, and stored at -20°C. All subsequent steps were performed at room temperature unless noted otherwise. On the day of staining, slides were warmed to room temperature, fixed in PBS + 4% PFA for 10’, and washed 3X in PBS. They were treated in 0.01M sodium citrate, pH 6 for 30’ at 75°C. They were then treated for 10’ in 0.2M HCl and 2’ in PBS + 1% Triton X-100. They were subsequently rinsed twice for 1’ each in PBS before a 10’ equilibration in 2X SSC + 10% formamide. Sections were exposed overnight at 37 °C to hybridization solution (10% dextran sulphate, 1mg/ml E. coli tRNA, 2mM Vanadyl Ribonucleoside, 200 μg/ml BSA, 2X SSC, 10% deionized formamide) containing 80nM oligo(dT)45 that was end-labelled with DIG Oligonucleotide Tailing Kit, 2nd generation (Roche) following the included instructions for short tails. On the following day, the slides were washed two times for 30’, each at 37 °C in 2X SSC + 10% formamide, and then rinsed briefly in 2X SSC followed by rinsing with PBST. Primary antibodies against digoxigenin (sheep polyclonal from Roche at 1:200) and FXR2P (A42; 1:500) were applied in blocking solution for 2 h. Tissue was rinsed before secondary antibody application in blocking solution for 1 h. Sections were rinsed and then mounted in mounting medium.

### Imaging

For quantification of FXG abundance, tissue sections were imaged through a 40X oil immersion objective (NA=1.15) using a Leica DMI4000 B inverted microscope coupled to a Hamamatsu Orca-R^2^ epifluorescent camera. For colocalization analyses, sections were imaged through the same objective using a Leica SPE II confocal microscope. Hippocampal images were acquired from the proximal CA3 region of the dorsal hippocampus. Cortical images were acquired dorsal to the anterior flexure of the corpus callosum.

### Quantification of FXG abundance

Epifluorescent micrographs of FXR2P immunostaining from the wild-type, *Fmr1* knockout, and I304N mice were processed to identify FXGs using previously published methods (Akins et al., 2012; Christie et al., 2009). In brief, images of entire sections were sharpened in Photoshop (Adobe) using the Unsharp Mask filter three times with the settings 500%, 2.5 pixels, and 128 levels. The identified puncta, which corresponded to FXGs, were counted. Rare puncta inappropriately identified as FXGs (e.g., autofluorescent blood vessels) were manually excluded from the quantifications. Statistical analyses were performed using GraphPad Prism 7.0e.

### Quantification of FXG composition

FXGs in confocal micrographs were identified based on FXR2P immunolocalization as described above. These FXGs were then manually annotated for the presence of signal in the FMRP channel blind to genotype. Statistical analyses were performed in R 3.5.1. Data were expressed as a binomial, with individual granules classified as FMRP-containing or FMRP-lacking. Genotype effects were analyzed using the glm function for each age and brain region with post-hoc pairwise comparisons between individual genotypes. Samples without variance (e.g., all FXGs lacking FMRP in the *Fmr1* nulls) prevented the model from converging. Therefore, a single *Fmr1* null data point was changed to FMRP-containing in all cases. Similarly, a single data point was changed to FMRP-lacking for P15 wild type cortex. These changes were conservative as they were in the opposite direction of the observed effect.

## Results

### The I304N mutation reduces FMRP association with FXGs in cortex and hippocampus

We first asked whether an intact KH2 domain is required for FMRP association with FXGs in cortical circuits. We investigated this possibility in brains from mice aged between postnatal days 15 and 30 (P15 to P30), the period during which FXGs are most broadly expressed in wild type mice as well as a period in which FXG abundance is regulated by FMRP (Christie et al., 2009). We immunostained brain sections from wild type, *Fmr1*^I304N^, and *Fmr1* null mice for FMRP and FXR2P. As in previous studies, we identified these FXGs using the FXR2P signal based on their distinctive morphology and location in deep layers of frontal cortex (Akins et al., 2012, 2017; Christie et al., 2009; Chyung et al., 2018). We then asked whether these cortical FXGs contained detectable FMRP signal (Fig. 1; Table 1). We detected FMRP in all FXGs in wild type mouse cortex, consistent with past studies (Chyung et al., 2018). Further, as expected, cortical FXGs in the *Fmr1* null brain did not contain detectable FMRP. In contrast to wild type FMRP, the mutant protein was detected in only 63% of FXGs at P15 and 31% of FXGs at P30. FMRP association with cortical FXGs is thus reduced in the *Fmr1*^I304N^ mice.

**Figure 1:**
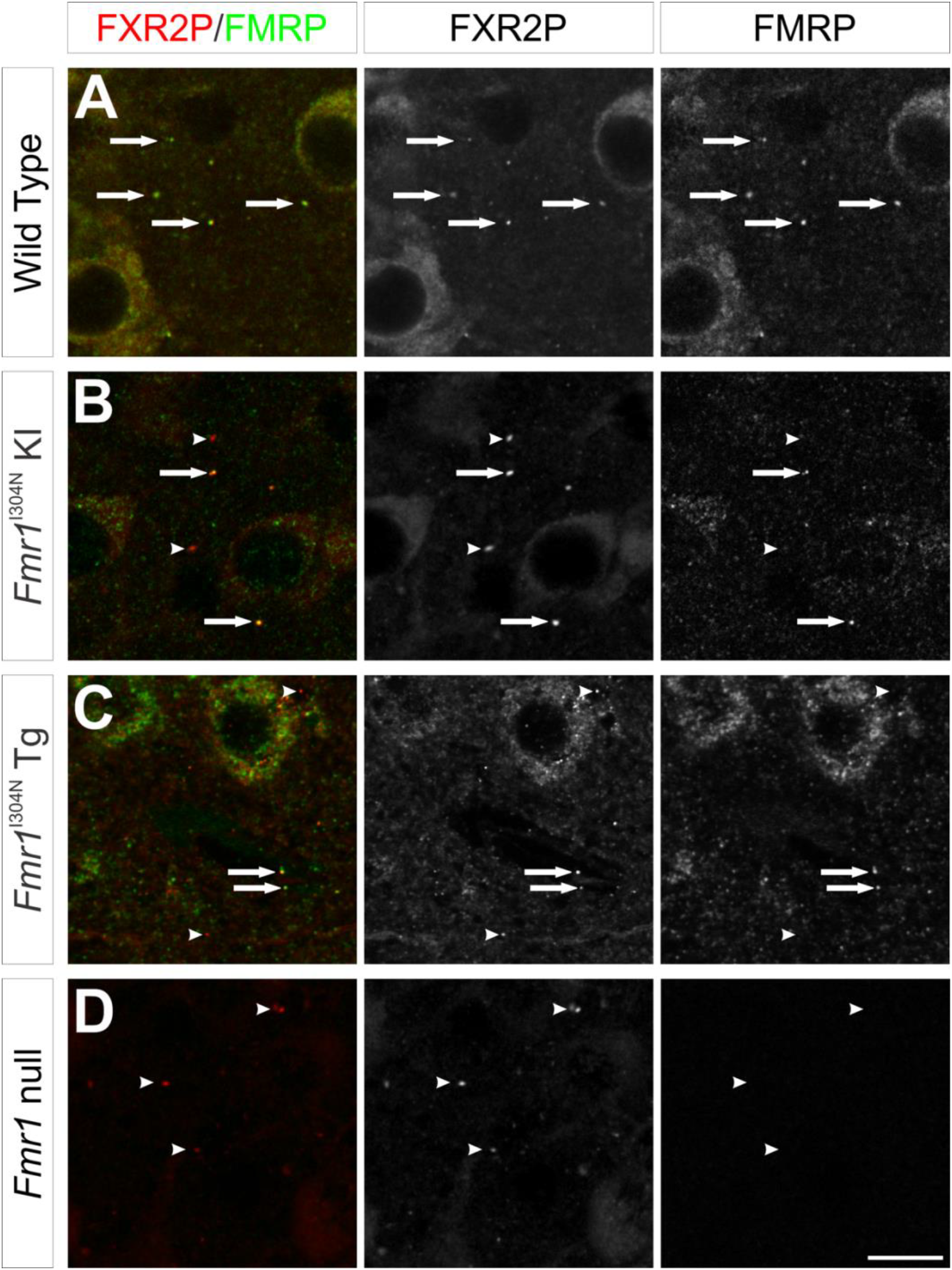
Genotype-dependent FMRP association with FXGs in cortex. Sections of P15 mouse cortex were stained for FXR2P (red) and FMRP (green). **(A)** In wild type tissue, FMRP was seen in all FXGs (arrows). **(B)** In *Fmr1*^I304N^ knockin tissue, the mutant FMRP colocalized with a subset of FXGs (arrows) but was absent from others (arrowheads). **(C)** In *Fmr1*^I304N^ transgenic tissue, the mutant FMRP colocalized with some FXGs (arrows) but not others (arrowheads). **(D)** In *Fmr1* null tissue, no FMRP signal was detected in FXGs (arrowheads). Scale bar = 10 μm.

**Table 1.**
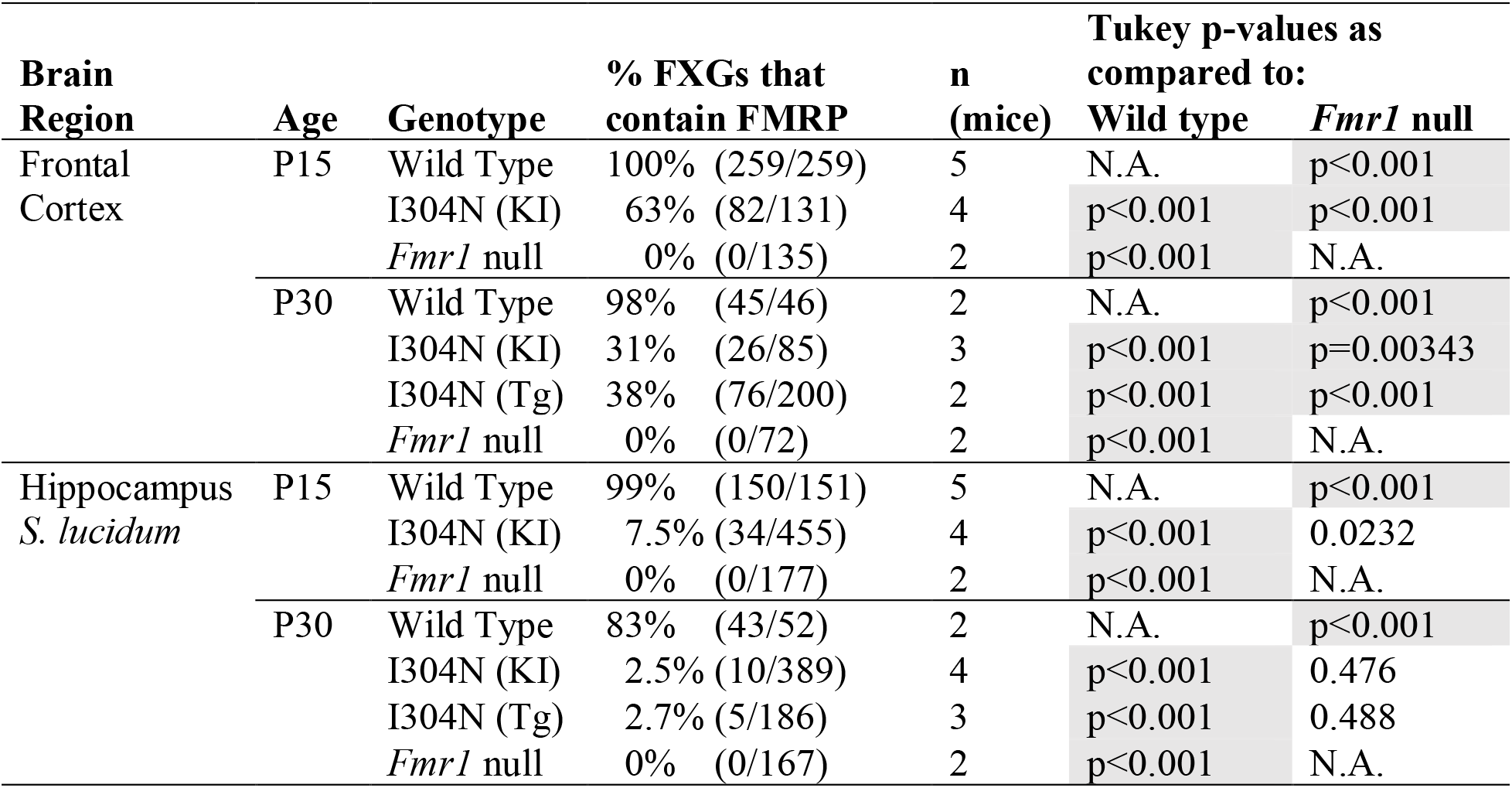
FMRP Content of FXGs. The portion of FXGs that contain FMRP in frontal cortex and stratum lucidum of hippocampal area CA3. For each brain region at each age, ANOVA revealed that genotype affected FXG composition (p<0.001). Post-hoc Tukey analyses were used to make pairwise comparisons between genotypes for each brain region and age. Comparisons that were significant with a threshold of 0.01 are indicated with shading. The two different lines (KI: knock-in and Tg: transgenic) of I304N mice did not detectably differ from each other in the P30 cortex (p=0.595) or hippocampus (p≈1). N.A.: not applicable.

We next addressed FMRP^I304N^ association with FXGs in hippocampal mossy fibers in stratum lucidum. Consistent with our past findings (Chyung et al., 2018), essentially all these FXGs in wild type mice contained FMRP while those in *Fmr1* null mice did not (Table 1; Fig. 2). In the *Fmr1*^I304N^ mice, only a small fraction (~8% at P15 and <3% at P30) of the FXGs contained the mutant FMRP^I304N^ protein. FMRP association with hippocampal FXGs is thus reduced in the *Fmr1*^I304N^ mice even more severely than with cortical FXGs.

**Figure 2:**
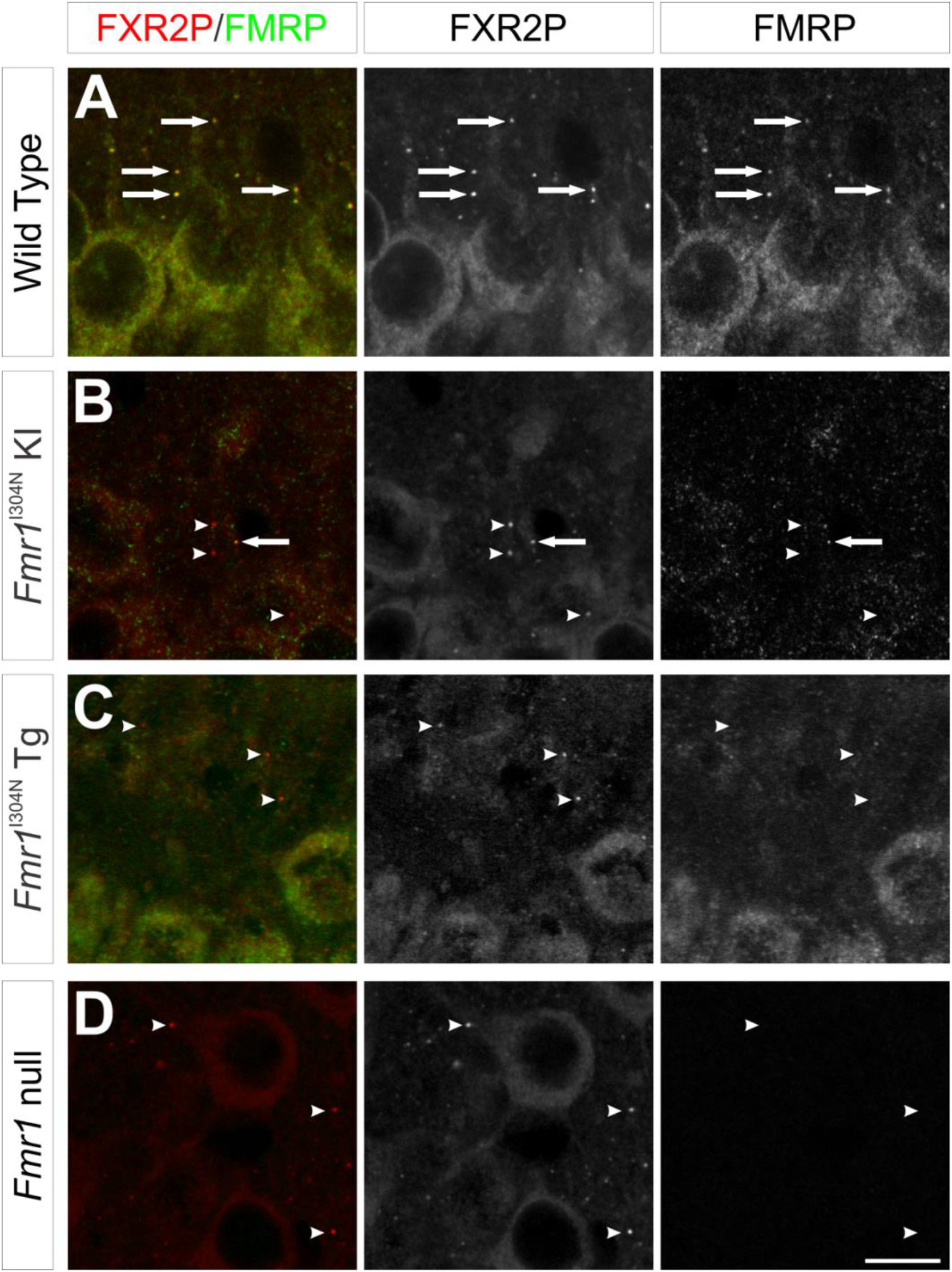
Genotype-dependent FMRP association with FXGs in hippocampus. Sections of P15 mouse hippocampus were stained for FXR2P (red) and FMRP (green) and FXGs identified based on FXR2P staining. **(A)** In wild type tissue, FMRP was seen in all FXGs (arrows). **(B)** In tissue from *Fmr1*^I304N^ knockin mice the mutant FMRP was absent from most FXGs (arrowheads) but colocalized with a small subset (arrows). **(C)** In *Fmr1*^I304N^ transgenic mice, the mutant FMRP was absent from nearly all FXGs (arrowheads). **(D)** In *Fmr1* null hippocampus, FMRP signal was not detected in FXGs (arrowheads). Scale bar = 10 μm.

We considered the possibility that the failure to detect FMRP^I304N^ associated with FXGs might be due to the lower expression of the mutant protein in the knock-in mice. To rule out this explanation, we examined tissue from mice that harbor a transgenic FMRP^I304N^ allele that is expressed at near wild type levels. As shown in Table I, the proportion of FXGs with associated FMRP^I304N^ was equivalent between the knock-in and transgenic models in both cortex (Fig. 1B-C) and hippocampus (Fig. 2B-C). The reduced association of FMRP^I304N^ with FXGs thus likely reflects an intrinsic property of this mutant protein rather than differential levels of expression. Together with the results above, these findings show that the KH2 domain is a key determinant of FMRP association with FXGs.

### I304N does not affect FXG association with ribosomes or mRNA

We next wanted to know whether loss of RNA and ribosome binding by FMRP affected FXG association with ribosomes and mRNA. Specifically, we wondered whether the presence of the mutant FMRP^I304N^ protein might disrupt these associations. Past studies have shown that ribosomes and mRNAs associate with FXGs in both wild type and *Fmr1* null brains (Akins et al., 2017; Chyung et al., 2018). We thus asked whether FXGs colocalized with ribosomes in *Fmr1*^I304N^ mice by immunostaining brain sections from all three genotypes for FXR2P, FMRP and 5S/5.8S rRNA (using the antibody Y10b). In particular, we focused our analyses on frontal cortex so that we could examine FXGs that contain the FMRP^I304N^ mutant protein. We found that both FMRP^I304N^-containing and FMRP^I304N^-lacking FXGs colocalized with ribosomes (Fig. 3A-B).

**Figure 3:**
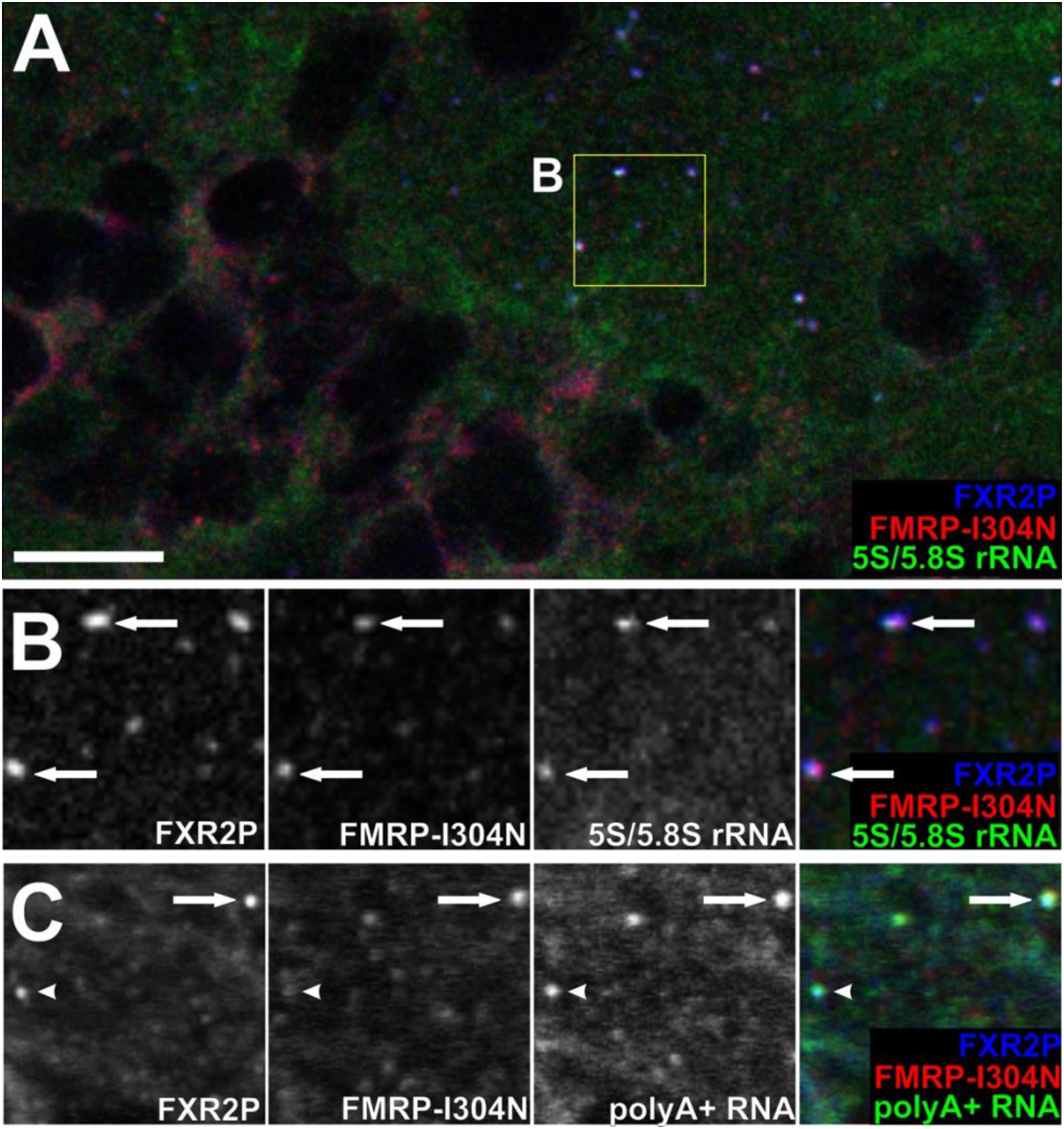
Ribosomes and mRNA are found in FMRP^I304N^-containing FXGs. Cortical sections from *Fmr1*^I304N^ mice were stained for FXR2P (blue) and FMRP (red) and FXGs identified based on the FXR2P signal. **(A,B)** In situ hybridization using oligo(dT) probes revealed polyA+ RNA in FXGs that contained the mutant FMRP (arrows) as well as those that did not contain detectable FMRP (arrowheads). **(C)** Immunostaining with the Y10b antibody revealed 5S and 5.8S ribosomal RNA (rRNA) in FMRP^I304N^-containing (arrows) and FMRPI304N-lacking (arrowheads) FXGs. Scale bar = 10 μm in A, 5 μm in B,C.

To test whether the mutant FMRP^I304N^ protein affected FXG association with polyA+ RNA, we combined immunostaining for FXR2P and FMRP with in situ hybridization with oligodT probes. As in both wild type and *Fmr1* null mice, FXGs across the *Fmr1*^I304N^ brain associated with polyA+ RNA (Fig. 3C), including FXGs that contained the mutant protein as well as those that lacked it. The I304N mutation in FMRP therefore did not detectably impact association of FXGs with RNA.

### I304N does not affect FXG substructure

We next asked whether FMRP^I304N^ impacts the organization of FXG components within the granules. To address this question, we used structured illumination super-resolution microscopy of CLARITY-treated sections from mice of all three genotypes (Fig. 4). We examined FXGs in cortex since these comprised a mixed population of FMRP^I304N^-containing and FMRP^I304N^- lacking granules in the mutant mice. Consistent with our past studies of hippocampal FXGs (Akins et al., 2017), we found that FMRP, FXR2P and rRNA in wild type cortical FXGs were arranged in overlapping but distinct domains. FXR2P and rRNA colocalized extensively within FXGs while FMRP colocalized with these proteins but was also found in a separate domain adjacent to FXR2P. This relationship was maintained in FXGs that contained the mutant FMRP^I304N^ protein. Similarly, FXR2P and rRNA colocalized within the granules in cortical FXGs that lacked FMRP, whether in the *Fmr1*^I304N^ or *Fmr1* null mice. Therefore, the presence of either wild type or mutant FMRP does not detectably alter the subgranular arrangement of FXR2P or rRNA.

**Figure 4:**
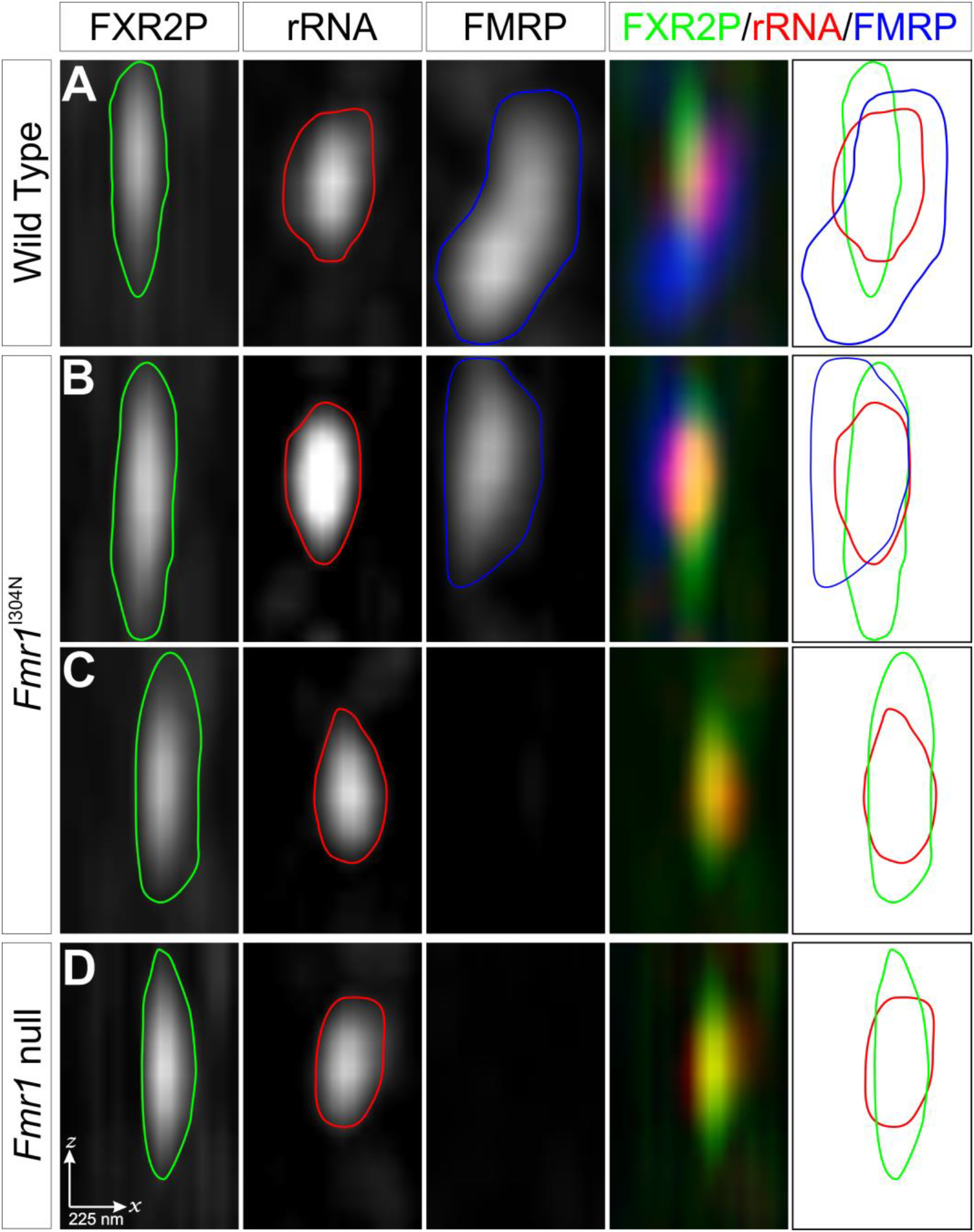
Substructure of FXGs. Cortical sections from P15 mice were stained for FXR2P (green), 5S/5.8S rRNA (red) and FMRP (blue) and imaged using structured illumination super-resolution microscopy. FXGs are depicted in the *xz* plane. In addition to single and merged immunofluorescence signals, traces of the signal in each individual channel are superimposed to show the subgranular structure. In FXGs that contained FMRP in **(A)** wild type and **(B)** *Fmr1*^I304N^ tissue, FXR2P, rRNA and FMRP were found in overlapping but distinct FXG domains. In FXGs that lacked FMRP, in both **(C)** *Fmr1*^I304N^ and **(D)** *Fmr1* null tissue, FXR2P and rRNA were found in an arrangement indistinguishable from the one they adopt in FMRP-containing FXGs. Scale bar = 225 nm.

### FMRP association with FXGs correlates with its regulation of their abundance

We next sought to interrogate the mechanisms whereby FMRP regulates FXG abundance by exploring whether intact KH2 functionality is required for this regulation. FXGs exhibit a developmental decrease in abundance that is regulated by FMRP (Akins et al., 2012, 2017; Christie et al., 2009). In the absence of FMRP there is an increase in their number. The I304N mutation impacts KH2-dependent functions cell wide but also limits the ability of FMRP to associate with FXGs. Thus, the I304N mutant mice allowed us to investigate whether FMRP regulation of FXG abundance reflects cell wide alterations in FMRP function or instead requires local FMRP function within the granules. Since the two I304N mouse lines exhibited equivalent FMRP association with FXGs, we conducted these studies in the knock-in line, in which the I304N mutation will impact all FMRP splice forms rather than just the one expressed in the transgenic line.

We first addressed this question in frontal cortex (Fig. 5). Consistent with our past studies (Christie et al., 2009), we found a significant increase in the number of FXGs in the *Fmr1* null cortex as compared to wild type at both P15 and P30. In cortex from *Fmr1*^I304N^ mice, we found an intermediate number of FXGs between that seen in wild type and *Fmr1* null. At P15, this number was significantly different from that seen in both wild type and *Fmr1* null mice. At P30, FXG abundance in *Fmr1*^I304N^ mice was not detectably different from either wild types or *Fmr1* nulls, despite these two genotypes differing significantly from each other, consistent with an intermediate phenotype in the *Fmr1*^I304N^ genotype.

**Figure 5:**
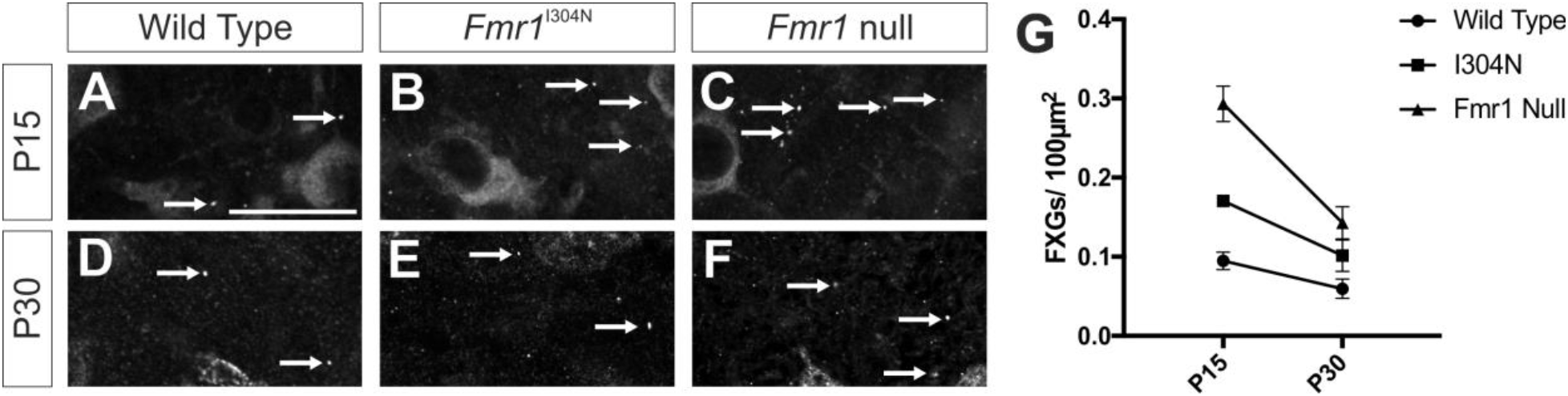
Genotype-dependent regulation of cortical FXG abundance across development. Cortical sections from **(A-C)** P15 and **(D-F)** P30 mice were immunostained for FXR2P and FXG abundance was determined in tissue from wild type (A,D), *Fmr1*^I304N^ knock-in (B,E) and *Fmr1* null (C,F) animals. A two-way ANOVA revealed significant effects of both genotype (p<0.0001) and age (p<0.0001) with a significant interaction between the two factors (p=0.0049). Post-hoc t-tests revealed that at P15, FXG abundance was significantly different between wild type and *Fmr1* null mice (p<0.0001). P15 *Fmr1*^I304N^ mice were different from both wild type (p=0.0054) and *Fmr1* null (p<0.0001) mice. At P30, wild type and *Fmr1* null mice differed significantly in their FXG abundance (p=0.0051), but *Fmr1*^I304N^ mice were not detectably different from either wild type (p=0.2110) or *Fmr1* null (p=0.2190) mice. n=6 P15 animals per genotype; n=5 P30 animals per genotype. Scale bar = 25 μm.

These findings raised the possibility that FMRP regulates FXG abundance locally within the granule, since a partial loss of FMRP from FXGs leads to an intermediate phenotype in *Fmr1*^I304N^ mice. We therefore asked whether loss of FMRP from nearly all FXGs, as seen in hippocampal mossy fibers, led to an increase in FXG abundance close to that seen in *Fmr1* null mice. Indeed, *Fmr1* null and *Fmr1*^I304N^ mossy fibers had equivalent FXG abundance to each other and significantly more FXGs than in wild type at both P15 and P30 (Fig. 6).

**Figure 6:**
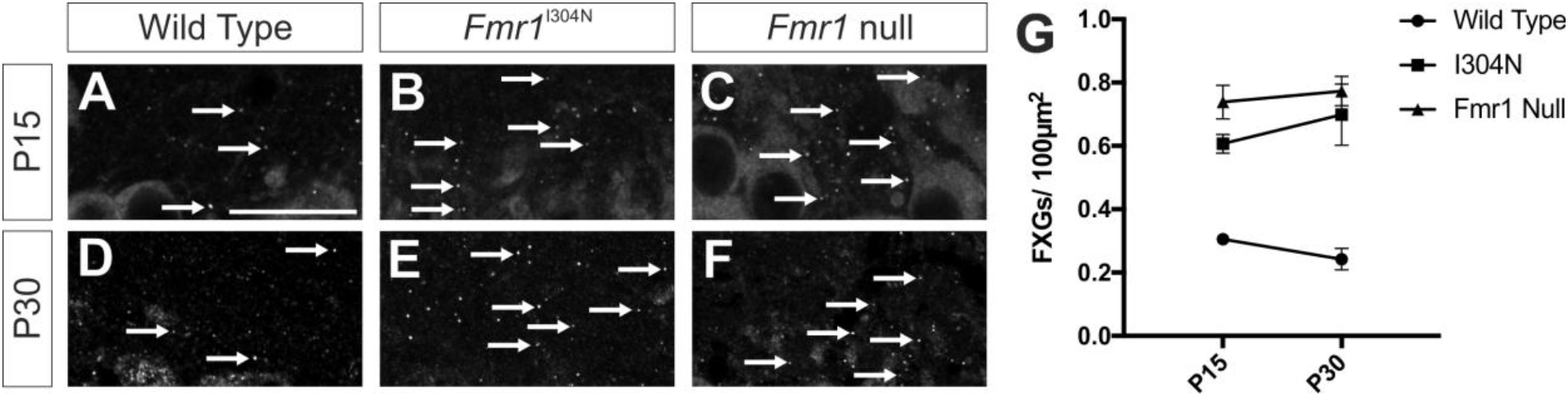
Genotype-dependent regulation of hippocampal FXG abundance across development. Hippocampal sections from **(A-C)** P15 and **(D-F)** P30 mice were immunostained for FXR2P and FXG abundance was determined in tissue from wild type (A,D), *Fmr1*^I304N^ knock-in (B,E) and *Fmr1* null (C,F) animals. A two-way ANOVA revealed a significant effect of genotype (p<0.0001) but not age (p=0.6093) with no significant interaction between the two factors (p=0.3132). Post-hoc t-tests revealed that at P15, FXG abundance was significantly different between wild type and *Fmr1* null (p<0.0001) and *Fmr1*^I304N^ (p=0.0004) hippocampus. No difference was detected between *Fmr1*^I304N^ and *Fmr1* null mice (p=0.1473). At P30, wild type mice differed from both the *Fmr1*^I304N^ and *Fmr1* null mice (p<0.0001 for both), while the mutant and null lines were indistinguishable (p=0.5812). n=6 P15 animals per genotype; n=5 P30 animals per genotype. Scale bar = 25 μm.

We next asked whether FXG abundance was related to the proportion of FXGs that contained FMRP. Since FXG abundance varies by age and brain region, we first normalized the FXG count for each condition to the wild type animals for that age and brain region. A correlation analysis revealed a negative relationship between FMRP content and FXG abundance (Fig. 7; p=0.001; Spearman ρ=-0.9203). Together, these findings indicate that FMRP regulation of FXG abundance requires the presence of the protein in FXGs but is independent of RNA binding by FMRP.

**Figure 7:**
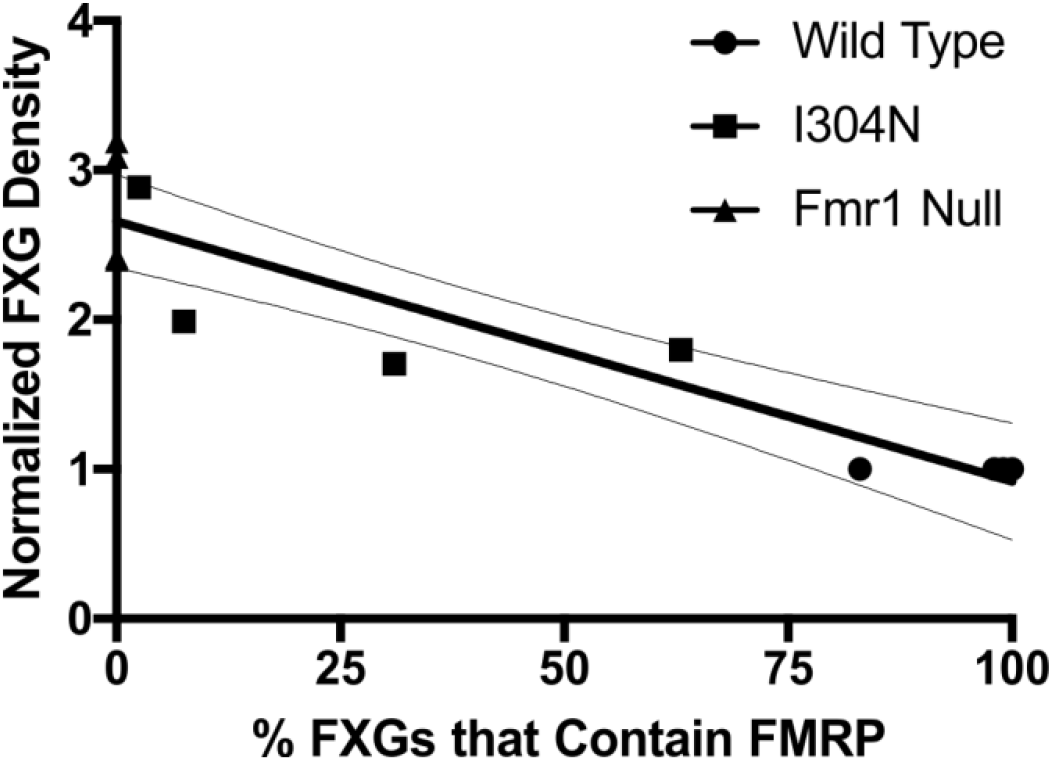
FXG abundance correlates with the percentage of FMRP-containing FXGs. FXG abundance for each age and brain region was normalized to wild type. The corresponding value was then correlated with the percent of FXGs for each condition that contain FMRP (Table 1). Spearman correlation revealed a significant negative correlation (p<0.0001; ρ = -0.9203), with abundance increasing as the number of FMRP-containing FXGs decreased. Depicted is a best fit linear regression along with 95% confidence intervals.

## Discussion

To elucidate mechanisms by which FMRP associates with FXGs and regulates their abundance, we asked how these granules were affected in *Fmr1*^I304N^ mice in which FMRP was present but could not directly bind RNA. This mutation had no obvious effect on ribosome or mRNA association with FXGs, consistent with these associations existing in the absence of FMRP (Akins et al., 2017). Instead, the ability of FMRP, which is normally found in essentially all forebrain FXGs, to associate with these granules was impaired. Interestingly, FMRP^I304N^ association with FXGs was brain region-dependent, with the mutant protein present in a sizable fraction of FXGs in frontal cortex but absent from nearly all FXGs in hippocampal mossy fibers. Super resolution imaging showed that both wild type and mutant FMRP exhibited similar distribution within the granules when present. Additionally, we found that FMRP regulation of the FXGs varied among brain regions in a way that depended on its presence in the granules.

FXGs thus illustrate the broad array of neuron type-specific mechanisms that regulate the formation, composition and subcellular distribution of FMRP-containing RNPs. Previous studies have demonstrated that neuron classes differ not only in whether or not they express FXGs but also in the protein and RNA composition of these granules. The current study reveals that FXGs in different classes of neurons assemble by distinct mechanisms. Together, these studies point toward a model in which upstream factors regulate the formation and axonal localization of FXGs in parallel to or downstream of the axonal localization of their target mRNAs. For example, FXGs localize to axons and associate with RNA and ribosomes independent of FMRP (Akins et al., 2017). Moreover, FXGs themselves are not required for the axonal transport of their target mRNAs (Akins et al., 2017). One likely possibility is that other RNA binding proteins beside FMRP control the assembly and targeting of axonal RNA granules and that FMRP associates with these granules in a manner that is heavily dependent on RNA binding.

The assembly of RNA granules is thought to occur in several conserved steps (Boeynaems et al., 2018; Buchan, 2014). These include reaching a critical local concentration of core RNA binding proteins which upon binding to each other or to mRNA undergo conformational changes that encourage fibrillization and granule formation (Burke et al., 2015; Elbaum-Garfinkle et al., 2015; Guo and Shorter, 2015; Kato et al., 2012; Molliex et al., 2015; Schwartz et al., 2013; Treeck et al., 2018; Weber and Brangwynne, 2012; Zhang et al., 2015). Secondary to this core formation is the further addition of peripheral shell proteins that interact with other cellular structures and can provide functionally distinct granule subdomains (Trcek et al., 2015; Weil et al., 2012; West et al., 2015; Wheeler et al., 2016). Although FXGs are a distinct family of RNA granules that differ from other granules including stress granules and P bodies (Christie et al., 2009), their assembly is likely governed by similar, conserved mechanisms. FXG core assembly appears driven by FXR2P, since it is a necessary FXG component and its association with ribosomes and mRNA in FXGs is unperturbed in the absence of FMRP (Akins et al., 2017; Christie et al., 2009). FMRP, in contrast, may be a peripheral FXG protein since the composition and structure of the granules are independent of FMRP while super resolution microscopy places this protein in a domain adjacent to that occupied by other FXG components. FMRP likely associates with the core components via protein-RNA and protein-protein binding mechanisms in parallel. To date, FXR and ribosome proteins are the only known FXG protein components. Identifying additional FXG components may elucidate the factors that control FXG assembly and/or transport. It will be particularly insightful if there are FXG core components that exhibit cell type-specificity in their expression that may explain why FXGs are only found in certain cells.

The restricted spatiotemporal pattern of FXG expression indicates that these granules are under tight regulation that varies depending on the brain region, neuronal cell type, and developmental stage. Consistent with this, FMRP association with FXGs is also heavily dependent on cellular context – FMRP^I304N^ associates with roughly half of FXGs in neocortical circuits but is only rarely found in hippocampal FXGs. Differential expression of FXG components that bind FMRP in a KH2-independent manner may explain such a difference. Notably, an intact KH2 domain is not required for FMRP to bind to FXR1P and FXR2P or to the FMRP interacting proteins 82-FIP, CYFIP1 or NUFIP (Bardoni et al., 2003; Laggerbauer et al., 2001; Linder et al., 2008; Ramos et al., 2006; Schenck et al., 2001; Zang et al., 2009). Among these, FXR1P is the leading candidate to underlie circuit-dependent differences in FMRP^I304N^ association with FXGs as FXR1P is a component of cortical FXGs but is not found in hippocampal mossy fiber FXGs (Chyung et al., 2018). Thus KH2-independent protein-protein interactions may stabilize FMRP association with FXGs either by providing a binding partner that helps anchor FMRP or by producing a change in FXG conformation that favors FMRP association (Fig. 8). Either one of these models raises the possibility that FMRP association with FXGs is dynamically regulated, potentially by signal-dependent posttranslational modifications. Phosphorylation may be a mechanism for regulating this association, as phosphorylation of the RNA binding protein Fus at residues in its low complexity domain can inhibit its assembly into higher order structures (Han et al., 2012; Murray et al., 2017). Phosphorylation of FMRP at low complexity domain residues that mediate granule association or at other sites that modulate translational regulation may provide multiple parallel mechanisms for FMRP to regulate axonal translation in a cell and context specific manner.

**Figure 8:**
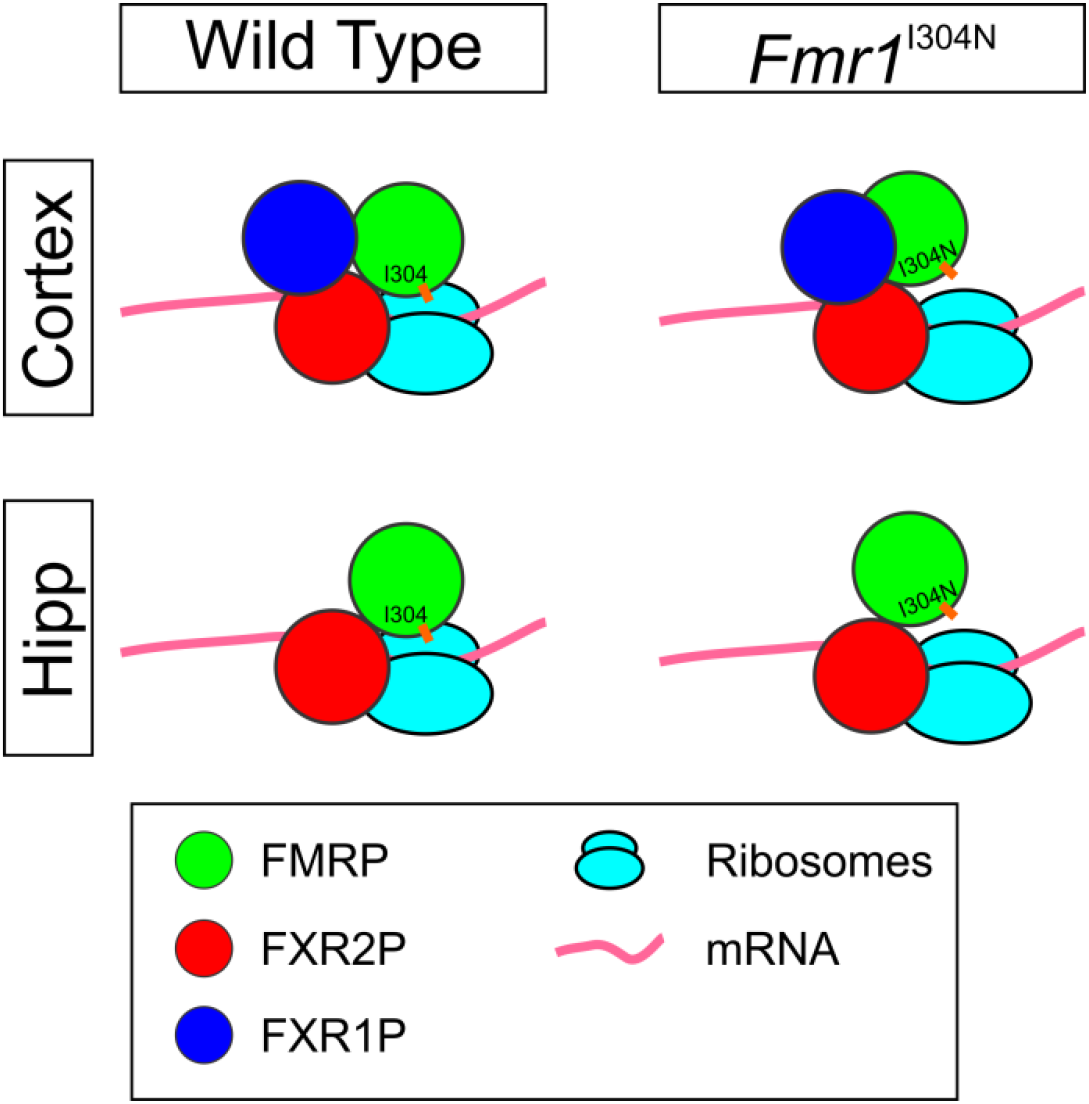
A model for I304-mediated FMRP association with FXGs. In wild type brains, FMRP associates with FXGs via I304-mediated protein-RNA interactions as well as protein-protein interactions mediated by other sites. When protein-RNA interactions are disrupted in the *Fmr1*^I304N^ mutant mice, FMRP association with FXGs is weakened. In cortical granules, FMRP binds to both FXR2P and other FXG components such as FXR1P, providing a stronger attachment than that found in hippocampal FXGs, which contain FXR2P but not FXR1P.

At least some of FMRP’s roles in FXGs do not involve translational control as our findings show that FMRP regulates FXG abundance via local, KH2-independent roles. In principle, FMRP could regulate cell-wide levels of a protein important for FXG formation or stability. Alternatively, FMRP could inhibit the local, FXG-dependent synthesis of a protein that in turn regulates FXG abundance. Our data do not support either of these possibilities since translational control by FMRP is not required to regulate FXG abundance. Instead, our data support a model in which FMRP regulation of FXG number requires local interactions that occur within the granules themselves. These interactions could involve the recruitment (or exclusion) of additional FXG component proteins that influence the propensity of FXG components to assemble into a granule. Alternatively, FMRP may compete with FXR2P for binding to other FXG components; the absence of FMRP may allow for a tighter association between FXR2P and these other components, thereby leading to a more stable and longer lived FXG. An additional possibility is that FMRP may be present in smaller RNPs where it inhibits their ability to coalesce into larger granules. Such a model implies that axonal FMRP-associated RNPs normally shuttle between FXGs and smaller complexes. Moreover, these smaller complexes would be depleted in *Fmr1* null axons as they are sequestered in FXGs. Either way, our findings raise the possibility that signals that regulate FMRP also regulate FXG dynamics.

Together, our studies of FXGs reveal that FMRP-containing RNPs are assembled via cell type-specific mechanisms resulting in distinctive regulatory protein and target mRNA composition. Moreover, these RNPs are differentially sensitive to mutations that affect FMRP structure and function. Patients with hypomorphic mutations, such as I304N, that impact FMRP function but are not complete nulls may thus exhibit tissue and cell type-dependent phenotypes that reflect the diversity of FMRP-containing RNPs. While I304N was the first FMRP point mutation discovered in a patient, and remains the most extensively studied, additional FMRP mutations have been found that result in typical or atypical FXS (Suhl and Warren, 2015). These mutations include one, R138Q, identified in a patient with mild FXS that disrupts FMRP binding to axonal potassium channels but spares RNA binding and translational control (Myrick et al., 2015). It will be informative, pending availability of an FMRP^R138Q^ mouse model, to determine whether this mutation, which impairs protein-protein interactions, results in region-specific alterations in the composition and function of FXGs or other FMRP-containing RNPs.

Finally, it seems likely that cell type-dependent formation and function of RNPs is a general phenomenon of which FXGs are a specific example and that other RNA binding proteins are also found in a variety of RNPs that vary among cell types. Mutations in these other RNA binding proteins may thus have cell type-dependent effects that reflect the diverse supermolecular complexes in which they are found. RNA binding protein dysfunction underlies a variety of nervous system disorders including FXS and autism but also amyotrophic lateral sclerosis, spinocerebellar ataxia, and spinal muscular atrophy (Conlon and Manley, 2017; King et al., 2012; Nussbacher et al., 2015). It will be of interest whether atypical or mild forms of these disorders arise from point mutations that produce cell type-dependent RNP phenotypes.

